# Geographical variation in plasma bactericidal ability of breeding tree swallows (*Tachycineta bicolor*)

**DOI:** 10.64898/2026.09.25.754392

**Authors:** David A. Chang van Oordt, Cedric Zimmer, Conor C. Taff, Daniel R. Ardia, Alexandra P. Rose, David A. Aborn, L. Scott Johnson, Thomas A. Ryan, Jennifer J. Uehling, Maren N. Vitousek

## Abstract

Immune defenses may vary across environments due to differences in life history or abiotic and biotic characteristics, but few studies have compared the strength of immune responses across broad geographical gradients. Here we investigated how bacteria killing ability (BKA), a measure of constitutive innate immune function, varies among populations of tree swallows (*Tachycineta bicolor*) inhabiting different environments. We hypothesized that populations living in unpredictable environments, which also have shorter breeding seasons and thus fewer opportunities to renest in the event of reproductive failure, would invest in reproduction at the expense of immunity. To test this hypothesis, we measured BKA and monitored reproductive activity in four populations located across most of the breeding range of tree swallows, in Tennessee, New York, Wyoming and Alaska. Consistent with our predictions, BKA was lowest in Alaska and Wyoming and highest in Tennessee. In most populations BKA was also lower during incubation than during nestling provisioning, and lower in males than in females. These results suggest that the high reproductive investment seen in populations experiencing more unpredictable breeding conditions may come at a cost to immune function. While reduced immune investment could increase susceptibility to parasites, this potential cost is likely offset by lower parasite pressure in these environments, limiting the need for maintaining strong constitutive innate defenses like BKA.

## Introduction

The ability of an organism to defend itself against pathogens—its immune defenses— can vary depending on the demands of its environment. On one side, spatial variation in parasite pressure, via differences in prevalence or biodiversity, may require increased investment in immune function. Accordingly, many studies find geographical variation in immune phenotypes and pathogen resistance and tolerance across different geographical scales and gradients in response to changing parasite exposure (Bonneaud et al. 2011; Vogelweith et al. 2013; Schmitt et al. 2017; Becker et al. 2020; Fecchio et al. 2020). On the other hand, differences in life history may need distinct resource allocation strategies that can also affect investment in immune function (McDade 2003; Stoks et al. 2006; Ardia 2007; Chang van Oordt et al. 2022).

Pathogen biodiversity, much like other biodiversity patterns, is expected to decrease at higher latitudes and elevations as abiotic conditions shift, including increasing environmental unpredictability where conditions like temperature or precipitation are highly variable (Rohde 1999; Willig et al. 2003; but see Preisser et al. 2022). Unpredictable environments also impose important selection pressures on host life history via lower host survival and shorter host breeding seasons, in turn favoring higher investment in each host reproductive attempt (Drent and Daan 1980; Stearns 1992). Considering that both reproductive and immune investment are energetically costly, immune function is often downregulated when resources are limited or during life stages with high energetic needs (French et al. 2007; Iseri and Klasing 2013; Ruoss et al. 2019). Thus, when there are trade-offs between investment in host reproduction and immune defense, more unpredictable environments should favor investment in reproduction at the expense of immune defense during breeding (Lochmiller and Deerenberg 2000; Norris and Evans 2000; Ardia 2005; Tieleman et al. 2005; French et al. 2007; Balasubramaniam and Rotenberry 2016). Both parasite biodiversity trends and reproductive trade-offs predict that immune defenses may be lower in organisms breeding in more unpredictable environments; but few studies have compared the strength of immune responses across unpredictability gradients (but see Ardia 2005, 2007).

Here we take a large-scale geographical approach to study whether environmental unpredictability, measured via temperature variability, can explain variation in immune investment across populations, and the degree to which reproductive-immune trade-offs impact immune phenotypes. We tested these relationships in tree swallows (*Tachycineta bicolor*), a widely-distributed migratory passerine with a broad breeding range spanning from the states of Georgia to Alaska in the United States, and throughout much of Canada (Winkler et al. 2020a). In tree swallows, breeding early is associated with higher lifetime fitness due to access to key resources like nesting sites and high-quality prey items (Twining et al. 2018; Winkler et al. 2020b). As latitude increases, so does the intensity of the negative association between breeding date and clutch size across the tree swallow range, suggesting that there is stronger selection on early breeding at high latitudes where breeding seasons are shorter than at lower latitudes (Winkler et al. 2014, 2020b; Taff et al. 2026). Similarly, at higher latitudes, nestling provisioning rates are elevated, suggesting higher reproductive investment (Alaska and Wyoming: Zimmer et al. 2020).

We hypothesized that tree swallow populations breeding in more unpredictable environments (typically those at high latitude and/or elevation) invest less in constitutive immune function because shorter breeding seasons in these environments favor individuals that spend more energy on reproduction. To test our hypothesis, we compared immune function among four breeding populations of tree swallows located in Tennessee, New York, Wyoming and Alaska in the United States of America (Figure 1B). Tennessee and Alaska are near the southern and northern edges of the breeding range of tree swallows, respectively. Meanwhile, the Wyoming and New York populations are at similar intermediate latitudes, but the Wyoming population is at higher elevation. Previous work in these populations has shown that birds in Alaska and Wyoming experience the shortest breeding seasons as well as the greatest temperature unpredictability during breeding—calculated as the standard deviation of the residuals of a daily ambient temperature additive model—while birds in Tennessee experience the longest breeding seasons and the lowest temperature unpredictability (Figure 1A, Zimmer et al. 2020).

**Figure 1.**
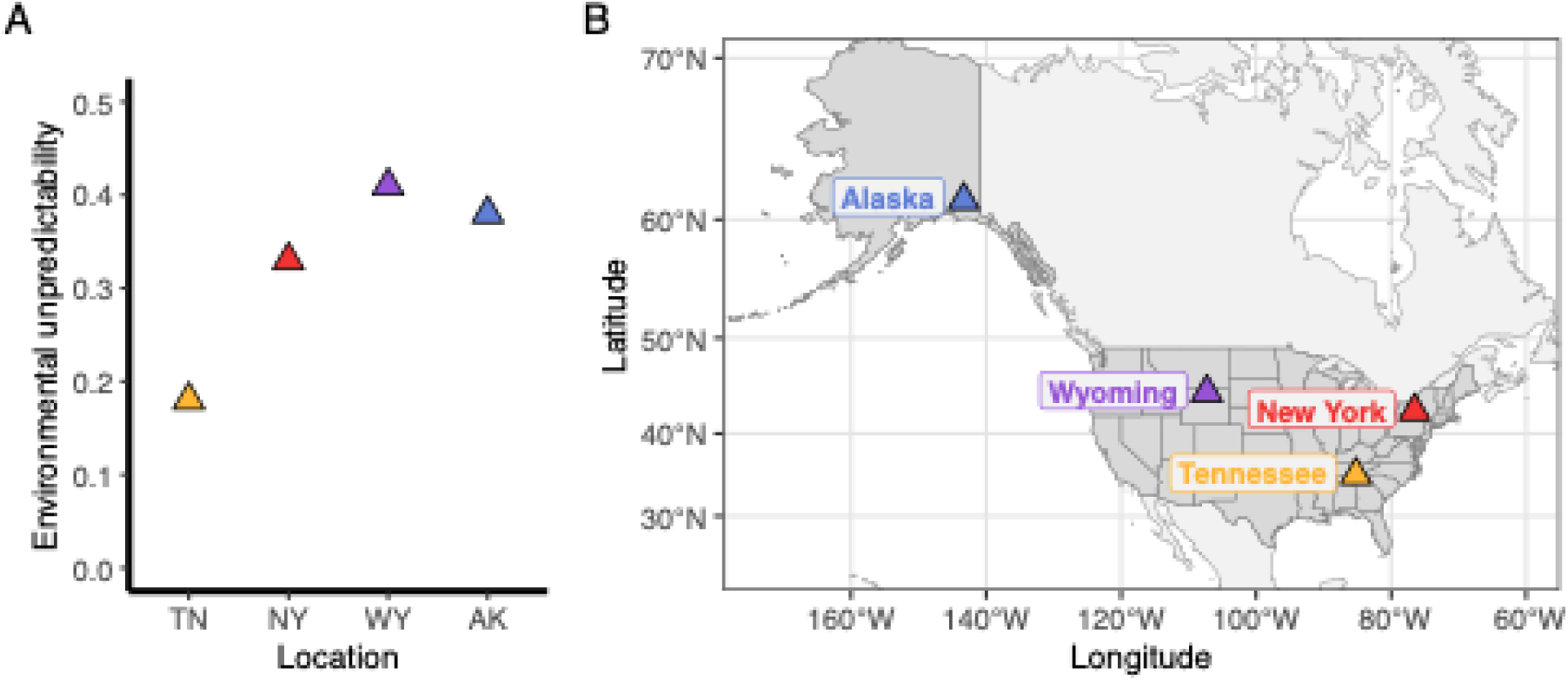
(**A**) Environmental unpredictability and (**B**) locations of study populations.

We measured bacteria killing ability (BKA), which is a functional metric that assesses the effective strength of the immune response by measuring constitutive innate immune function (Matson et al. 2006; Millet et al. 2007). The assay is performed *ex vivo*, which allows measuring standing activity of the complement system and circulating antimicrobial peptides (Millet et al. 2007). We predicted that BKA would be the lowest in Alaska and Wyoming, where temperatures are more unpredictable, and highest in Tennessee, where environmental temperatures are most predictable.

In contrast to induced responses—those activated after exposure to a pathogen— constitutive immune responses can show the state of standing immunity and how environmental conditions may constrain investment; however, to our knowledge, no study has assessed constitutive innate immune function across large geographical gradients (but see Schmitt et al. 2017 for landscape scale effects). Previous work has shown that tree swallows breeding in harsher, more northerly environments have weaker inflammatory responses to subcutaneous phytohemagglutinin injections, although antibody production against sheep red blood cells was the same in all populations (Ardia 2007). Moreover, more northern populations experience stronger trade-offs between immunity and reproduction (Ardia 2005). Within populations, female tree swallows mount weaker inflammatory responses when they have large brood sizes, and this relationship is more negative at higher latitudes (Ardia 2005). Additionally, tree swallows in high latitude populations trade off inflammation with antibody production, but their counterparts in lower latitudes can simultaneously mount strong responses in both arms of the immune system (Ardia 2007).

We also predicted that, across populations, immune investment decreases between incubation and provisioning because the reduced likelihood of renesting in the event of a nest failure during provisioning would select for greater investment in reproduction than immunity, and nestling provisioning requires a large energy budget that can trade off against immune investment (Chang van Oordt et al. 2022). We also predicted that this difference would be more pronounced in less predictable environments and/or those with short breeding seasons because short breeding seasons are more likely to favor parental investment at the expense of immunity (Ardia 2005). Finally, we predicted that, within populations, BKA would be higher in earlier breeding birds because early breeders can have more access to resources and may be able to mount strong immune responses while maintaining high reproductive investment due to higher individual quality (Verhulst and Nilsson 2008; Winkler et al. 2020b; Chang van Oordt et al. 2022).

## Materials and Methods

### Study populations and field samples

We used frozen blood plasma samples that had been collected from male and female tree swallows in 2017 and 2018 and previously used for the study by Zimmer et al. (2020). Tree swallows were caught and sampled in Chattanooga, Tennessee (51.1°N, 85.2°W, 206 m, 2018); Ithaca, New York (42.5°N, 76.5°W, 340 m, 2018); Burgess Junction, Wyoming (44.5°N, 107.3°W, 2451 m, 2018); and McCarthy, Alaska (61.4°N, 143.3°W, 445 m, 2017).

Zimmer et al. (2020) previously showed that thermal unpredictability differs among these populations using the standard deviation of the residuals of a time series model of temperature (Franch-Gras et al. 2017). Alaska and Wyoming have the highest standard deviations, indicating more unpredictability, Tennessee had the lowest dispersion indicating less unpredictability, and New York had intermediate unpredictability.

Sample and data collection consisted of monitoring nest boxes to obtain nesting data and capture individuals (see Zimmer et al. 2020). We recorded the onset of laying (‘lay date’), clutch size, hatching date, brood size, average mass of nestlings on the sixth day after hatching, fledging success and the number of nestlings that fledged successfully from each nesting attempt. Zimmer et al. (2020) previously reported that clutch size did not vary between these populations, nor did fledging success—except for the Wyoming population where fledging success was lower, likely as a result of unusually cold conditions during breeding in the years of study. Female tree swallows were caught twice in different stages of the nesting attempt: during incubation (six to eight days after all eggs were laid), and during the nestling feeding stage (six to eight days after eggs hatched). Male tree swallows were captured 6–12 days after eggs hatched. All birds were weighed and sampled for blood and received an individually numbered USGS aluminum band.

The blood samples were taken in sets of three: one sample within three minutes of capture, a second sample 30 minutes after capture before they received an injection of dexamethasone, a synthetic glucocorticoid (to test rapid negative feedback in the HPA axis), and the third sample 30 minutes after the injection (Zimmer et al. 2020; Taff et al. 2023).

Blood samples were collected in the field, and we separated plasma less than 4 hours after collection. The separated plasma samples were stored at -20 °C in the field and then placed in a freezer at -80 °C when they arrived at the lab. The samples were thawed to analyze steroid concentrations for different studies (Zimmer et al. 2020; Taff et al. 2023), and then refrozen at -80 °C until this investigation. To salvage the largest possible sample size for this study, we pooled all three plasma samples from a capture event into a single sample per individual and breeding stage to increase the amount of plasma available for the immune assay. Given that samples in a sequence were taken within a short time frame (approximately one hour between first and last sample), we do not expect significant differences in BKA as the time would not be sufficient to ramp up production of complement and antimicrobial proteins. After pooling, the final number of samples analyzed for each location, sex and breeding stage varied between 17 and 71 plasma samples (Table 1).

**Table 1.**
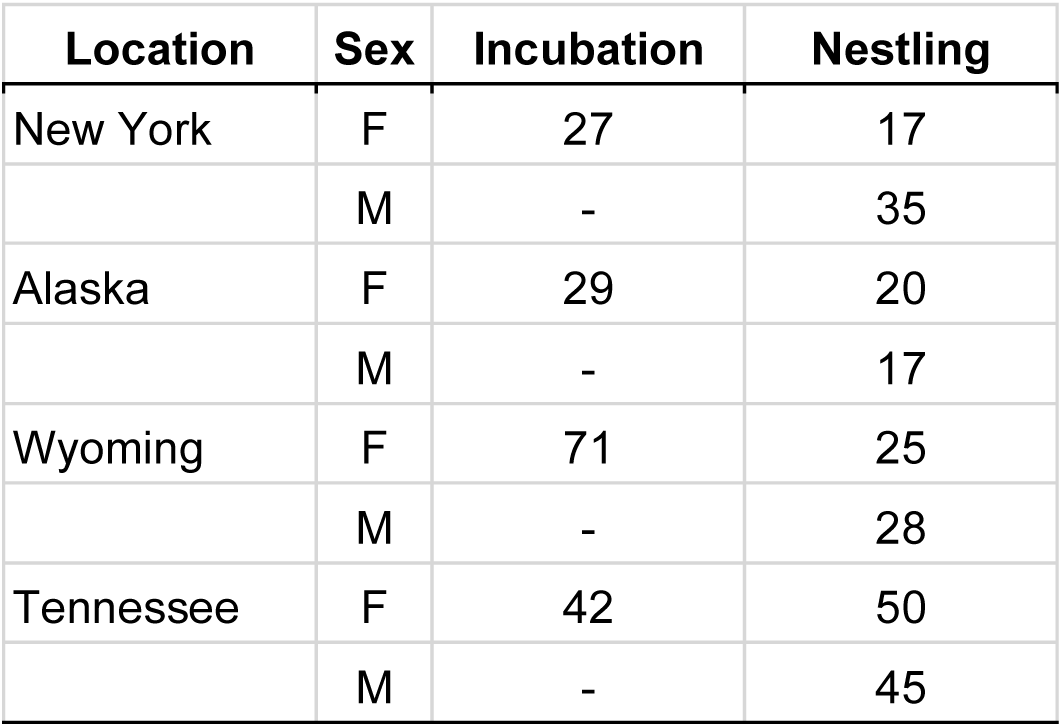
Sample size by location, sex and nesting stage.

| Location | Sex | Incubation | Nestling |
| --- | --- | --- | --- |
| New York | F | 27 | 17 |
|  | M | - | 35 |
| Alaska | F | 29 | 20 |
|  | M | - | 17 |
| Wyoming | F | 71 | 25 |
|  | M | - | 28 |
| Tennessee | F | 42 | 50 |
|  | M | - | 45 |

### Bacteria Killing Assay

We measured BKA (Matson et al. 2006; Millet et al. 2007) from the plasma samples by doing bacteria killing assays using a protocol modified from French and Neumann-Lee (French and Neuman-Lee 2012) for 96-well microplates. First, we reconstituted a pellet of *Escherichia coli* ATCC® 8739 (Microbiologics Epower 0483E7™) in 40 mL of PBS prewarmed to 37 °C, then we vortexed and incubated the suspension in a water bath at 37 °C for 30 minutes. We diluted the stock solution to a working concentration of 10^5^ CFU mL^-1^. Neither the stock solution nor the working solution was kept for more than 24 hours. We diluted 5 µL of plasma in 13 µL of PBS in each well and added 6 µL of the working bacterial suspension. The mixture was then incubated at 41 °C for 30 minutes alongside a blank and a positive control consisting of 24 µL of PBS and 18 µL of PBS with 6 µL of the working bacterial suspension, respectively. After the incubation step at 41 °C, we added 125 µL of cold Tryptic Soy Broth to each well and measured the absorbance of the mixture at 300 nm using a Synergy HTX Multi-mode reader (Agilent Technologies Inc., Santa Clara, California, USA). We then incubated the plate for 12 hours at 37 °C. After incubation, we resuspended the bacteria in the wells using a microplate shaker at 800 rpm for at least two minutes and measured absorbance again at 300 nm after the final incubation step.

We calculated BKA for each well using the following equation:

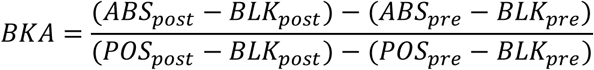

Where ABS represents the sample absorbance, BLK is the average of all the blank controls and POS is the average of all positive controls. Meanwhile, ‘pre’ and ‘post’ subindices indicate whether the absorbance measurement was taken before or after incubating at 37 °C, respectively. The adjustment using blank absorbance was used to correct for changes in TSB absorbance after incubation, and the use of ‘pre’ and ‘post’ incubation measurements accounted for variation in among-individual sample absorbance caused by different plasma hues (Hill 1995). We ran each sample in triplicate when enough plasma was available. Because we took advantage of the remaining samples from another study that had gone through multiple freeze-thaw cycles, most values were very small and assay variability was elevated (mean within-sample coefficient of variation: 149.02%). Despite the elevated dispersion, we used the entire BKA dataset because most samples were exposed to similar freeze-thaw conditions and we still expect broad patterns to arise, but we limit our interpretation to population level trends.

### Statistical analysis

We compared BKA and the relationship between BKA and life history using linear and generalized linear mixed methods using the package ‘lme4’ in the statistical language R version 4.2.1 (Bates et al. 2015; R Core Team 2024). We then used the package ‘emmeans’ to make *post hoc* Tukey’s multiple comparison tests between locations (Lenth 2023).

We tested whether lay date and body mass differed between populations to complement the life history comparisons reported by Zimmer et al. (2020). We used linear regression models to assess differences in lay date and body mass across populations. The body mass model included location and sex and used male and female data from the nestling provisioning stage only, while the lay date model included only location using female data as a reference for the entire nest.

Then, we tested the predictors of variation in BKA using linear mixed models. Since we ran each sample up to three times, we included individual ID as a random factor. We compared BKA between locations using the whole dataset with a general model that included lay date, location, sex and breeding stage as fixed variables, and individual ID as a random factor. Finally, we tested the location-specific effect of breeding stage (incubation vs. nestling provisioning) on BKA using female samples only. This model included lay date, location, and breeding stage as fixed factors, and individual ID as a random factor. We included an interaction between location and breeding stage to test whether populations differ in the degree to which BKA is suppressed during the provisioning period.

## Results

Some life history traits varied between locations. Average lay date among birds included in this study was different among all locations during the years of study (Tennessee: May 2nd ± 1.0 day S.E.; New York: May 20th ± 1.2; Alaska: June 4th ± 1.2; Wyoming: June 10th ± 0.8; all pairwise comparisons: *p* < 0.001, Supplementary Materials Table S1–2). Males had lower body mass than females during nestling provisioning (*p* < 0.001); and body mass was also lower in Wyoming (*p* < 0.001) compared to the other three populations (Supplementary Materials Table S3).

BKA also differed between populations (Figure 2A, Supplementary Materials Table S4). It was highest in Tennessee (mean = 0.35 ± 0.06), and lowest in Alaska (mean = 0.05 ± 0.06) and Wyoming (mean = 0.03 ± 0.05). BKA was also lower for males than for females during the provisioning period (Figure 2B, Supplementary Materials Table S4). Contrary to our prediction, among females, BKA was significantly lower during incubation than during provisioning in Alaska, Wyoming, and Tennessee (Figure 2C, Table 2). The degree of upregulation of BKA during provisioning was similar across Tennessee, Wyoming and Alaska, but greater than in New York (Table 2).

**Figure 2.**
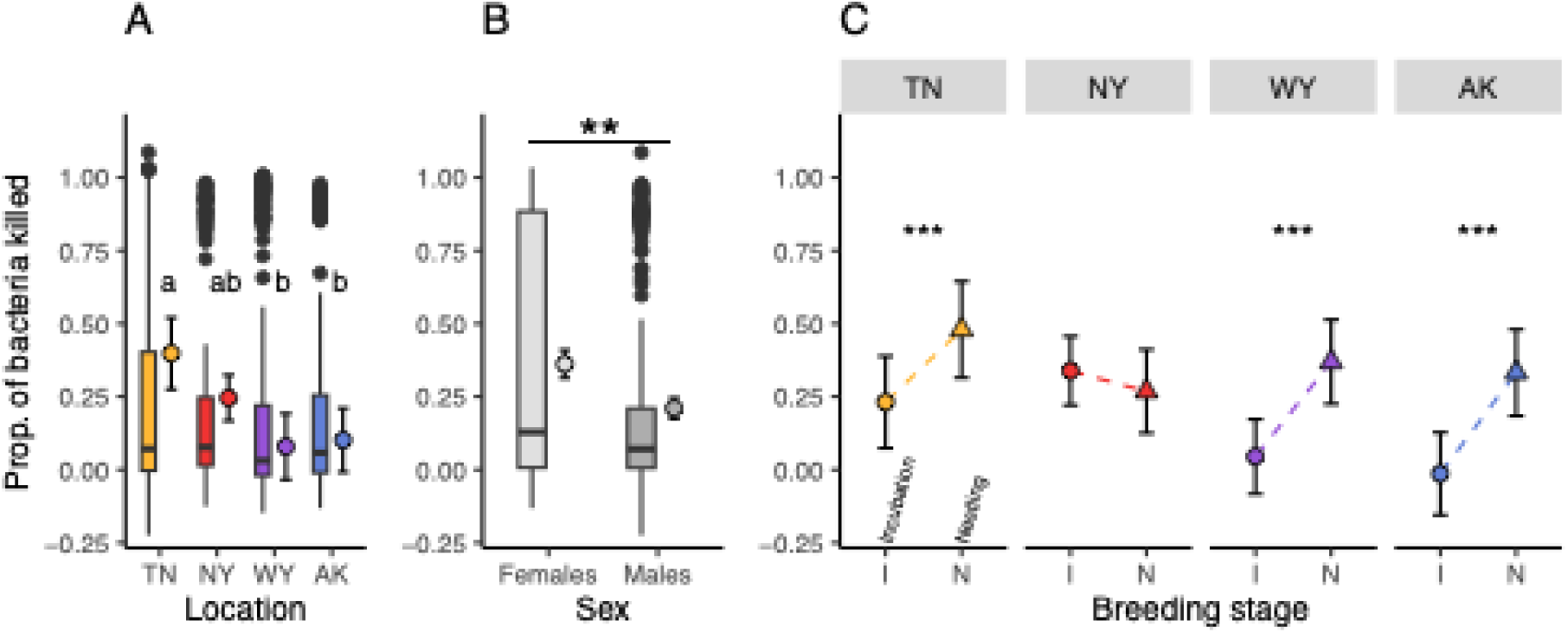
Bacterial killing ability (BKA) at each location separated by location (**A**), sex (**B**) and life history stage in adult females (**C**). In **A** and **B**, points alongside the boxplots show the mean BKA for each location (**A**) or sex (**B**), along with the 95% confidence interval of the mean as calculated from *post hoc* analysis. In **C**, points show the mean BKA of female tree swallows during incubation (circles) and the nestling provisioning stage (triangles) and the 95% confidence interval of the mean based on *post hoc* analysis. Dashed lines show the direction of change between life history stages. Letters in **A** group locations that are not statistically different from each other based on post hoc pairwise comparisons. In **B** and **C**, two asterisks (**) indicate *p* < 0.01, and three asterisks (***) indicate *p* < 0.001.

**Table 2.** Summary of the mixed model BKA including an interaction between breeding stage and location. This model uses data from female tree swallows only. The table includes the coefficient estimates, their confidence intervals, and *p*-values.

| **Predictors** | **Estimates** | **CI** | **p** |
| --- | --- | --- | --- |
| (Intercept) | -0.16 | -0.97 – 0.66 | 0.706 |
| Lay date | 0 | -0.00 – 0.01 | 0.239 |
| Location [Alaska] | -0.35 | -0.54 – -0.16 | **<0.001\*\*\*** |
| Location [Wyoming] | -0.29 | -0.48 – -0.11 | **0.002\*\*** |
| Location [Tennessee] | -0.11 | -0.29 – 0.07 | 0.246 |
| Nesting Stage [Nestling] | -0.07 | -0.20 – 0.06 | 0.281 |
| Location [Alaska] * Nesting Stage [Nestling] | 0.42 | 0.26 – 0.58 | **<0.001\*\*\*** |
| Location [Wyoming] * Nesting Stage [Nestling] | 0.39 | 0.24 – 0.55 | **<0.001\*\*\*** |
| Location [Tennessee] * Nesting Stage [Nestling] | 0.32 | 0.17 – 0.47 | **<0.001\*\*\*** |

## Discussion

Our study highlights how host susceptibility can vary across broad geographical scales and contributes to a growing body of evidence on macroecological patterns of host immune defenses (Martin et al. 2004; Ardia 2005, 2007; Arriero et al. 2015). BKA was lower in populations that breed in more unpredictable thermal environments (Figure 2A), suggesting that immunity tracks other biogeographical patterns. Male tree swallows also exhibited lower BKA than females, even though both males and females provide care during the nestling provisioning period (Leffelaar and Robertson 1986; Whittingham et al. 2003). It is unclear why males and females differ in their bactericidal capacity as they are seemingly experiencing similar constraints, but intrinsic differences between male and female physiology may be the cause. For example, immune suppression could occur from elevated testosterone in males (Duffy et al. 2000; Lipshutz and Rosvall 2021), or changes in luteinizing hormone and prolactin in females after incubation (Mota-Rojas et al. 2023). Shifting endocrine profiles as the breeding season moves forward may allow females to upregulate BKA and recover from a period of increased disease susceptibility to levels that males may not reach due to testosterone-associated immunosuppression.

The geographical patterns of BKA indicate that tree swallow populations in more unpredictable environments may be more vulnerable to infections given their lower investment in immunity. However, in these environments, reduced immune investment may not be costly if fewer parasites and pathogens are present. As parasite pressure decreases, so do the benefits of maintaining constitutive immune defenses. Therefore, it may be beneficial to lower investment in immunity and avoid immunopathology when stronger immune defenses are not necessary (Råberg and Stjernman 2003; Graham et al. 2005).

While we did not quantify parasite abundance, the observed differences in immune function among populations could be seen even in the absence of differences in parasite pressure if populations inhabiting more unpredictable environments with shorter breeding seasons invest more in reproduction at the expense of immunity. These processes could also be operating together if the release of parasite pressure in more unpredictable environments favors resource allocation strategies with higher investment in reproduction than in immune defenses. In this case, low parasite exposure in unpredictable environments may help offset some of the potential costs associated with high reproductive investment and low immune investment.

If parasite diversity and exposure do not decrease with higher unpredictability, it is possible that the environmental conditions that lead to higher reproductive investment and shorter breeding seasons also shape immune function, suggesting unpredictable environments impose similar selection pressures on life history and innate immunity. It is unclear how parasite pressure effectively changes with more unpredictable environments. Parasitic worm biodiversity and abundance increase at higher latitudes (Johnson and Haas 2021), but changes in complement-dependent immune mechanisms such as BKA are ineffective against those parasites. The complement system is more effective against microbes, but there is no evidence of a latitudinal reduction in bacterial diversity in tree swallow nests (Forsman 2016). Thus, the effect of changing parasite diversity abundance on host immunity relative to the pressures imposed by life history remain unknown.

Our results also show that immune investment can differ within a short period of time as female tree swallows transition from one breeding stage to another. Contrary to our predictions, BKA was lowest during the incubation period, and increased during the nestling provisioning stage. One possible explanation for this pattern is that pathogen pressure may increase over the course of the breeding season, and thus elevated immune function would be increasingly necessary (Merino et al. 2000; Cosgrove et al. 2008). Moreover, female tree swallow behavior between incubation and nestling provisioning changes sharply, which may lead to a significant shift in pathogen exposure. Incubating females spend most of their time in the nest, while provisioning females spend more time foraging to provide for their offspring. Extended time outside of the nest may increase exposure through increased interactions with the environment or conspecifics, especially considering that tree swallows are more likely to visit nests other than their own during this period (Taff et al. 2021). Another possible explanation is that incubation may be more energy-demanding than nestling provisioning, so it exhibits greater trade-offs with immunity. Studies in other taxa find that immune function often declines during the more costly reproductive stages. However, the most common pattern in birds is that nestling provisioning is more energetically costly than incubation (Nord and Williams 2015), and evidence suggests that tree swallows are likely no different (Williams 1988; Boyle et al. 2012; Chang van Oordt et al. 2022).

The change (or lack thereof) in BKA between incubation and nestling provisioning also differed for the New York population compared to the other three populations (Figure 2C, Supplementary Materials Table S5). The coefficient estimates of the interaction terms showed a similar significant upregulation in BKA between incubation and provisioning in Alaska, Wyoming and Tennessee; but not in New York. The apparent lack of differences in BKA across nesting stages in New York could result from smaller sample sizes (Table 1–2); or it could reflect a real difference in investment in immune function across reproductive stages, potentially resulting from other unmeasured differences in environment across populations at mid-latitudes. Alternatively, given that we used archived samples from a previous experiment, the samples left over for our analyses may have been biased towards individuals that started nesting later in the season which were not included in other experiments. Median lay date in the preceding years was approximately May 13^th^ (Shipley et al. 2020) while the mean lay date for the birds included in our BKA analysis was May 20^th^.

Individuals that lay later in the season are generally considered to be low quality individuals with lower lifetime reproductive success (Winkler et al. 2020b), and are subject to stronger negative associations between BKA and other traits like clutch size or baseline glucocorticoids (Chang van Oordt et al. 2022; Chang van Oordt et al. 2024). Thus, late breeders may exhibit different immune phenotypes than their earlier-breeding conspecifics.

Our results demonstrate that although bactericidal assays are sensitive to freezing (Jacobs and Fair 2016; Claunch et al. 2022), it is nevertheless possible to uncover broad immunological patterns from plasma samples frozen for four to five years. Bactericidal potency can decrease during freezing, but plasma proteins are less susceptible to freezing than immune cells, albeit they may be susceptible to consecutive freeze-thaw cycles. By showing that we can still recover patterns, our study may open opportunities for even broader comparative studies that incorporate multiple years and populations.

Any future studies should consider the complexity of the immune system in trying to understand these phenotypes under natural conditions. Immune processes differ in specificity, potency, energy requirements and effectiveness, and thus the collective immune phenotype may differ from a single metric of immune function. In general, we predict a similar pattern of regulation across all costly components of the immune system. The cost of investing in immune responses may increase as environmental temperature unpredictability increases, and the opportunities to reproduce decrease. Thus, we would expect immune responses with higher costs to be downregulated under high temperature unpredictability and reduced reproductive opportunities. Meanwhile, other parts of the immune system (those with lower associated costs) may show different patterns of regulation across environments if some populations can still afford investment in those responses. Our study provides insight into the regulation of complement and antimicrobial proteins, which are part of a first-line-of-defense immune mechanism and are constitutively expressed in the bloodstream. A previous comparative study, also in tree swallows, found that the inflammatory response to a novel antigen, which can be costly (Murtaugh et al. 1996; Iseri and Klasing 2013), shows similar patterns as those in our study, decreasing with latitude (Ardia 2007). In contrast, antibody production to a novel antigen, an indicator of the adaptive immune response which can be less costly than the inflammatory response (Iseri and Klasing 2013), did not differ across locations (Ardia 2007), showing that low cost immune responses may be energetically affordable even when environmental conditions are less favorable. Thus, our result shows a similar pattern to that of an inflammatory response, but an immune trait that is less energetically costly or does not carry the same risk of immunopathology may not show variation along an environmental gradient if all populations can afford to invest more heavily in this particular mechanism. Our results also suggest that these geographical patterns in immune investment are not primarily driven by latitude, but by environmental conditions, as birds in a lower latitude but high elevation population (Wyoming), with a similar thermal unpredictability to a higher latitude population (Alaska), showed similar patterns of immune regulation.

Broad geographical approaches like those used in this study can advance our understanding of environmental and life history constraints on immunity and the limitations on disease spread and transmission. Overall, our findings suggest geographical variation in immune function, as birds in unpredictable environments invested less in a low-cost component of immune function. The degree to which these differences confer more or less protection against pathogens, as well as the degree to which differences in resistance provide a selective advantage in unpredictable environments, remains unclear. However, it is evident that variation in environment and life history may impose important constraints on the immune phenotypes of tree swallows across their range.

## Acknowledgments

This project would not have been possible without the field crew members who collected samples across all sites over the years, including Allison Anker, Abigail Blackstone, Jeremy Collison, Stephanie Cook, Alex Dopkin, Audrey Fox, Tamara Egans Harper, Brianna Johnson, Christine Kallenberg, Raisa Kochmaruk, Ann Li, Pascale Lubbe, Laura Marsh, Sarah Mirza, Calum Poulin, Alyssa Rodriguez, Yosvany Rodriguez, Kwame Tannis; and special thanks to Danica Lee, who designed the starting protocol for the microwell-based bactericidal assay; and Ava Ciaccia, Gracey Brouillard and Audrey Su, for their help in sorting and pooling samples and running the assays.

## Declarations

### Funding

Funding was provided by the Cornell Lab of Ornithology to DCV, and by the National Science Foundation (U.S.A.) IOS-1457251 and DARPA D17AP00033 to MV.

### Conflicts of interest

The authors declare to have no conflicts of interest.

### Ethical approval

All applicable institutional, state and national guidelines for the care and use of animals were followed, and all work was conducted under the appropriate IACUC, state, and federal permits.

### Data availability

Data will be made available in Figshare upon acceptance.

### Author contributions

DCV conceived the question and wrote the manuscript along with MV. DCV also performed the immunoassays and analyzed the data. All authors collected data in the field and provided feedback on the manuscript.

## Notes

### Competing Interest Statement

The authors have declared no competing interest.

