## Supplementary Tables for "Geographical variation in plasma bactericidal ability of breeding tree swallows (*Tachycineta bicolor*)"

Supplementary Materials

**Table S1**

| ***Predictors*** | ***Estimates*** | ***CI*** | ***p*** |
| --- | --- | --- | --- |
| (Intercept) | 139.08 | 136.64 – 141.52 | **<0.001** |
| Location [Alaska] | 15.26 | 11.94 – 18.59 | **<0.001** |
| Location [Wyoming] | 21.16 | 18.28 – 24.04 | **<0.001** |
| Location [Tennessee] | -17.71 | -20.85 – -14.57 | **<0.001** |
| Observations | 155 | | |
| R2 / R2 adjusted | 0.872 / 0.869 | | |
| **Table S1.** Summary of the model comparing lay dates between locations. '(Intercept)' refers to the New York population in Ithaca, NY. | | | |

### Table S2

| **contrast** | **estimate** | **SE** | **df** | **t.ratio** | **p.value** |
| --- | --- | --- | --- | --- | --- |
| Ithaca - Alaska | -15.26 | 1.68 | 151 | -9.06 | **<0.001** |
| Ithaca - Wyoming | -21.16 | 1.46 | 151 | -14.50 | **<0.001** |
| Ithaca - Tennessee | 17.71 | 1.59 | 151 | 11.14 | **<0.001** |
| Alaska - Wyoming | -5.89 | 1.39 | 151 | -4.25 | **<0.001** |
| Alaska - Tennessee | 32.98 | 1.52 | 151 | 21.67 | **<0.001** |
| Wyoming - Tennessee | 38.87 | 1.27 | 151 | 30.66 | **<0.001** |
| **Table S2.** Tukey's pairwise comparison test for lay date (in day of year) between locations where 'estimate' denotes the magnitude of the difference between groups. | | | | | |

### Table S3

| ***Predictors*** | ***Estimates*** | ***CI*** | ***p*** |
| --- | --- | --- | --- |
| (Intercept) | 18.73 | 18.33 – 19.13 | **<0.001** |
| Location [Alaska] | 0.21 | -0.32 – 0.74 | 0.437 |
| Location [Wyoming] | -0.91 | -1.38 – -0.44 | **<0.001** |
| Location [Tennessee] | 0.25 | -0.17 – 0.68 | 0.239 |
| Sex [M] | 0.91 | 0.59 – 1.22 | **<0.001** |
| Observations | 217 | | |
| R2 / R2 adjusted | 0.239 / 0.225 | | |
| **Table S4**. Model summary for body mass by location and sex. | | | |

### Table S4

| ***Predictors*** | ***Estimates*** | ***CI*** | ***p*** |
| --- | --- | --- | --- |
| (Intercept) | -0.83 | -1.42 – -0.23 | 0.006 |
| Lay date | 0.01 | 0.00 – 0.01 | **0.001** |
| Location [Alaska] | -0.16 | -0.30 – -0.02 | **0.024** |
| Location [Wyoming] | -0.18 | -0.32 – -0.04 | **0.010** |
| Location [Tennessee] | 0.14 | 0.02 – 0.26 | **0.027** |
| Breeding Stage [Provisioning] | 0.25 | 0.21 – 0.30 | **<0.001** |
| Sex [M] | -0.18 | -0.25 – 0.10 | **<0.001** |
| **Random Effects** | | | |
| σ2 | 0.03 | | |
| τ00 Individual_Band | 0.01 | | |
| ICC | 0.8 | | |
| N Individual_Band | 331 | | |
| Observations | 1042 | | |
| Marginal R2 / Conditional R2 | 0.021 / 0.800 | | |
| **Table S4**. Summary of the model testing the effect of location on BKA. The table includes the coefficient estimates, their confidence intervals and *p*-values. Below, the table shows statistics for the random effects: sample ID and individual ID. | | | |

**Table S5**

| **contrast** | **estimate** | **SE** | **df** | **t.ratio** | **p.value** |
| --- | --- | --- | --- | --- | --- |
| Incubation New York - Provisioning New York | 0.07 | 0.07 | 690.88 | 1.08 | 0.961 |
| Incubation New York - Incubation Alaska | 0.35 | 0.10 | 258.62 | 3.62 | **0.008** |
| Incubation New York - Provisioning Alaska | 0.01 | 0.10 | 286.60 | 0.06 | 1.000 |
| Incubation New York - Incubation Wyoming | 0.29 | 0.09 | 289.42 | 3.12 | **0.042** |
| Incubation New York - Provisioning Wyoming | -0.03 | 0.10 | 343.09 | -0.30 | 1.000 |
| Incubation New York - Incubation Tennessee | 0.11 | 0.09 | 274.10 | 1.16 | 0.943 |
| Incubation New York - Provisioning Tennessee | -0.14 | 0.10 | 313.26 | -1.48 | 0.819 |
| Provisioning New York - Incubation Alaska | 0.28 | 0.11 | 310.92 | 2.57 | 0.171 |
| Provisioning New York - Provisioning Alaska | -0.06 | 0.11 | 336.87 | -0.57 | 0.999 |
| Provisioning New York - Incubation Wyoming | 0.22 | 0.11 | 342.61 | 2.05 | 0.447 |
| Provisioning New York - Provisioning Wyoming | -0.10 | 0.11 | 386.42 | -0.88 | 0.987 |
| Provisioning New York - Incubation Tennessee | 0.04 | 0.09 | 341.08 | 0.39 | 1.000 |
| Provisioning New York - Provisioning Tennessee | -0.21 | 0.10 | 375.45 | -2.20 | 0.352 |
| Incubation Alaska - Provisioning Alaska | -0.35 | 0.05 | 567.28 | -7.12 | **< 0.001** |
| Incubation Alaska - Incubation Wyoming | -0.06 | 0.07 | 232.12 | -0.79 | 0.994 |
| Incubation Alaska - Provisioning Wyoming | -0.38 | 0.08 | 300.07 | -4.71 | **< 0.001** |
| Incubation Alaska - Incubation Tennessee | -0.24 | 0.13 | 284.51 | -1.91 | 0.545 |
| Incubation Alaska - Provisioning Tennessee | -0.49 | 0.13 | 324.74 | -3.73 | **0.006** |
| Provisioning Alaska - Incubation Wyoming | 0.29 | 0.08 | 259.90 | 3.74 | **0.005** |
| Provisioning Alaska - Provisioning Wyoming | -0.04 | 0.08 | 326.01 | -0.44 | 1.000 |
| Provisioning Alaska - Incubation Tennessee | 0.10 | 0.13 | 307.23 | 0.77 | 0.994 |
| Provisioning Alaska - Provisioning Tennessee | -0.15 | 0.14 | 348.75 | -1.09 | 0.958 |
| Incubation Wyoming - Provisioning Wyoming | -0.32 | 0.05 | 662.16 | -7.20 | **< 0.001** |
| Incubation Wyoming - Incubation Tennessee | -0.19 | 0.13 | 306.61 | -1.40 | 0.855 |
| Incubation Wyoming - Provisioning Tennessee | -0.43 | 0.14 | 352.21 | -3.17 | 0.035 |
| Provisioning Wyoming - Incubation Tennessee | 0.14 | 0.14 | 338.99 | 1.00 | 0.974 |
| Provisioning Wyoming - Provisioning Tennessee | -0.11 | 0.14 | 384.49 | -0.78 | 0.994 |
| Incubation Tennessee - Provisioning Tennessee | -0.25 | 0.04 | 612.75 | -6.47 | **< 0.001** |
| **Table S5**. Pairwise comparison of BKA by nesting stage and location. The table includes the estimated difference in BKA, their standard errors, degrees of freedom, *t*-ratio and *p*-values. | | | | | |
